# An interpretable alphabet for local protein structure search based on amino acid neighborhoods

**DOI:** 10.1101/2025.04.21.649886

**Authors:** Saba Zerefa, Jesse Cool, Pramesh Singh, Samantha Petti

## Abstract

**Motivation:** Recent advancements in protein structure prediction methods have vastly increased the size of databases of protein structures, necessitating fast methods for protein structure comparison. Search methods that find structurally similar proteins can be applied to find remote homologs, study the functional relationships among proteins, and aid in protein engineering tasks. The structure comparison method Foldseek represents each protein structure as a sequence of “3Di” characters and uses highly optimized sequence comparison software to search with this alphabet. An alternate alphabet encoding richer features has the potential to improve search accuracy while leaving the underlying search algorithm unchanged.

**Results:** We design a “3Dn” structural alphabet that encodes the local neighborhoods around each amino acid in an interpretable way. In a search benchmark task, a combination of our alphabet and Foldseek’s 3Di alphabet, outperforms each alphabet individually and ranks best among local search methods that do not require amino acid identity information. We provide software tools that enable the exploration of novel alphabets and combinations of alphabets for protein structure search.

**Availability and implementation:** The code is freely available at https://github.com/spetti/structure_comparison.

## 1 Introduction

Recent developments in protein structure prediction methods have resulted in dramatic increases in the size of protein databases, with AlphaFold alone providing predicted structures for each of the over 250 million proteins in UniProt as of 2025 [15] [4]. The newly expansive size of protein databases demands fast and computationally efficient algorithms for high quality protein structure analysis. Search methods that find structurally similar proteins can be applied to find remote homologs, study the functional relationships among proteins, and aid in protein engineering tasks. Structural alignment algorithms like TM-align and Dali effectively compare protein structures based on their three dimensional configurations; however, these algorithms are too slow and computationally expensive for searching large databases [24, 12].

In contrast, classic sequence comparison methods utilize dynamic programming [1, 6, 9] to efficiently search databases with hundreds of millions of proteins. Traditionally, these methods only compare primary sequences. The authors of Foldseek [23] designed an alternate alphabet that encodes information about the three-dimensional structure of the protein. By using this alphabet within established highly optimized search software designed for amino acid sequences, they created a tool for structure comparison search that is tractable for large databases without a large drop in accuracy as compared to Dali and TM-Align [12, 24].

Here we focus on designing an alternate structural alphabet for structure comparison search. Foldseek uses a neural network with a VQVAE architecture to learn an alphabet of twenty 3Di characters. The 3Di characters encode the relationship between an amino acid and its nearest amino acid neighbor nonadjacent in sequence. Due to the black-box nature of the machine learning components involved in training, the 3Di characters are difficult to interpret, making it challenging to extract biochemical meaning of any given 3Di character. Additionally, only using the single nearest neighbor to an amino acid when constructing 3Di characters limits the amount of structural information captured by the 3Di characters. The nearest neighbor method may be sensitive when comparing two closely related proteins if there are multiple amino acids at a similar distance. An alphabet encoding richer features has the potential to improve the accuracy of the search, while leaving the remainder of the search algorithm unchanged. Thus, the construction of alternate alphabets merits further investigation.

We develop interpretable methods that encode the local structure around an amino acid by taking into account the locations and secondary structure of nonadjacent neighbors within a fixed radius around the amino acid. By considering all neighbors within a fixed radius, we are able to capture local structure information beyond just a single neighbor, and by focusing on interpretable methods, we are able to extract biochemical meaning from our alphabet characters. We compare our alphabet, which we call 3Dn (*n* for “neighborhood”) to Foldseek’s 3Di alphabet on a protein search benchmarking task. We also compare combinations of these alphabets with the amino acid alphabet with BLOSUM62 scoring and an alphabet derived from backbone dihedral angles. Alphabet combinations that do not require amino acid identities may be of particular interest for searching when homology is not expected, e.g. for protein engineering applications or in cases of convergent evolution.

### 1.1 Summary of contributions

Our results can be summarized as follows:

- As a precursor to our “3Dn” alphabet, we first introduce *blurry neighborhoods* to describe the locations of nonadjacent neighbors of an amino acid within 15Å. We devise an accompanying Jaccard metric to score the similarity of two blurry neighborhoods. Using this scoring scheme, we achieve significantly improved performance on a structure search task as compared to search with Foldseek’s 3Di alphabet, and comparable results with the combination 3Di and amino acid alphabet.
- We introduce the 3Dn alphabet, built by clustering the blurry neighborhoods. The 3Dn alphabet performs marginally worse than the 3Di alphabet on a protein database search task, but combining the 3Dn and 3Di alphabets improves on the performance of each individual alphabet. The combined 3Di-3Dn alphabet has comparable performance to the slower blurry neighborhood approach and is the current state of the art alphabet combination for local search that doesn’t require amino acid identity information.
- We visualize the neighborhoods encoded by each 3Dn character. Then, we analyze the co-occurrence of 3Di and 3Dn characters with each other, amino acid identity, and an alphabet of backbone dihedral angles built by a mutual information clustering algorithm. We conclude that the 3Dn and 3Di alphabets are capturing different aspects of the conserved structure, thus explaining the improved performance of the combined alphabet.
- We establish a framework for exploring new structural alphabets. In addition to an efficient implementation of the search benchmarking task, we provide code to train BLOSUM-like matrices, combine multiple alphabets, and reduce alphabet size via clustering based on mutual information. The software is independent of the type of structure information used, and thus can be used with other structural alphabets.

### 1.2 Related work

Protein structure comparison methods experience tradeoffs with respect to algorithm efficacy, computational efficiency, and interpretability. Algorithms like TM-align and Dali are accurate [24] [12], yet the computational expense of such techniques makes them infeasible for large protein database searching tasks. Works such as AlphaFind [20], PLMSearch (protein language model) [16], and DHR (dense homolog retriever) [13] utilize deep learning techniques to construct an embedding for the whole protein and infer similarity by comparing embeddings. While these methods allow for efficient global search, they are not effective for local search in which only parts of the protein are conserved.

Here we focus on developing an alphabet that can be used with existing local search tools. We compare against the aforementioned Foldseek 3Di alphabet, which was trained with a VQVAE architecture. Recent work trained an alternate alphabet and accompanying scoring scheme in an end-to-end manner using message-passing architecture of ProtMPNN [5]. In contrast, we specifically focus on deriving an alphabet from explicitly defined features to arrive at an interpretable, black-box free alphabet.

Works such as [10, 14] train neural networks to predict the Foldseek 3Di characters from the protein primary sequence, thus enabling structural search without access to a predicted structure. The same methods could be applied to predict any alphabet or combination of alphabets from the primary sequence.

## 2 Methods

### 2.1 Representing local amino acid neighborhoods

We quantify similarity between a pair of amino acids in different structures by comparing the locations and secondary structures of the amino acids in close proximity to them. We consider two amino acids of the same sequence to be *neighbors* if their alpha carbons are within 15Å of each other, and they are more than 5 positions away in the sequence.

First we designed a common reference frame to compare the relative positions of the neighbors between two proteins. The reference frame is defined with respect to an individual amino acid and is independent of rotation or translation of the greater protein. The reference frame of an amino acid satisfies three rules: the alpha carbon lies at the origin, the vector between the alpha carbon and beta carbon lies on the positive *z* axis^1^, and the nitrogen lies on the *XZ* plane, with positive *X* coordinate. We discretize the sphere with radius 15Å into 250 equal sized sections using spherical coordinates (see Appendix A.1). We assign each neighbor to one of 1000 “bins” determined by which of the 250 sections of the sphere the neighbor occupies and the secondary structure of the neighbor: beta sheet, left helix, right helix, or other, as determined by the backbone dihedral angles (see Appendix A.1).

For each amino acid, we define an *n-hot vector* 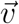 encoding the locations and secondary structure of neighbors: 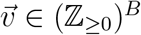 where *B* is the number of bins. The value of 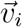 represents the number of neighbors of amino acid *x* that are assigned bin *i*.

#### 2.1.1 Blurry neighborhoods

Our goal is for amino acids with similar *n*-hot vectors to be assigned the same structural character. Although *n*-hot vectors capture the local structure around an amino acid, their discrete nature poses challenges in comparing two different *n*-hot vectors. For instance, between two structurally similar proteins, the bin of an analogous neighbor may not be completely conserved but rather moved to an adjacent bin. To address this, we introduce the blurry neighborhood, which is a continuous vector that builds upon the notion of the *n*-hot vector by capturing patterns of how neighborhoods differ between structurally similar proteins. By working in the continuous space of blurry neighborhoods as opposed to the discrete space of *n*-hot vectors, we can more meaningfully compare the local structure between two amino acids.

The blurry neighborhood of an amino acid is the expected distribution over bins that its neighbors would occupy after one “evolutionary timestep.” Our training set of aligned structures contains pairs of proteins in the same family, superfamily, or fold; we do not assume that the aligned pairs are homologous (e.g. they may be similar due to convergent evolution rather than a shared common ancestor). However, for simplicity in computing our blurry neighborhoods we consider each pair aligned structures in our training set to be one “evolutionary time step” apart.

We generate blurry neighborhoods with a learned transition matrix that encodes the probabilities of neighbors moving between bins. Thus, the blurry neighborhood intrinsically incorporates information regarding the relatedness of particular bins. The construction of a blurry neighborhood is a multi-step process, which is outlined as follows.

Define matrix *C ∈* (ℤ_*≥*0_)^(*B*+1)*×*(*B*+1)^ where *C*_*ij*_ represents how often a neighbor in one structure is in bin *i* and the analogous neighbor in the aligned structure is in bin *j*. Formally, let *C*_*ij*_ be the number of times we observe aligned pairs of positions *x*_*a*_, *x*_*b*_ in protein *X* and *y*_*a*_, *y*_*b*_ in protein *Y* (across all training structural pairwise alignments) in which *x*_*a*_ is aligned to *y*_*a*_, *x*_*b*_ is aligned to *y*_*b*_, *x*_*b*_ is in bin *i* of the discretized sphere around *x*_*a*_, and *y*_*b*_ is in bin *j* of the discretized sphere around *y*_*a*_. If *y*_*b*_ is not within 15Å of *y*_*b*_, this pair contributes to the count *C*_*i,−*1_, where negative one represents an extra bin indicating an amino acid that is not within 15Å. Since it is not meaningful which protein serves as *X* and which as *Y* within each pair, we symmetrize by setting *F* = (*C* + *C*^*⊺*^)*/*2.

We construct transition matrix *T ∈ ℝ* ^(*B*+1)*×*(*B*+1)^ by assigning 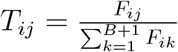. Thus, *T*_*ij*_ represents the probability of a neighbor moving from bin *i* to bin *j* after one step of evolution. The blurry neighborhood 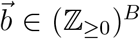 represents the expected distribution of neighbors of an amino acid after an evolutionary timestep. We compute the blurry vector 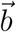 from the *n*-hot vector 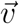 as follows. First we append the value 4 to 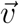 to represent neighbors in the negative one bin and call this (*B* + 1)-dimensional vector 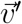.This accounts for the possibility that there are neighbors outside the 15Å radius that may move in and yields the nice property that applying the transition matrix does not affect the population-wide average number of neighbors. Then we obtain 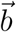 by computing 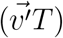 and ignoring the position corresponding to the negative one bin.

### 2.2 Comparing local amino acid neighborhoods with the Jaccard metric

We utilize the generalized Jaccard metric to compare the similarity between the blurry neighborhoods corresponding to two amino acids from different proteins. Let 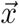 and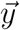 represent the blurry neighborhoods associated with amino acids *x* and *y*, respectively. The Jaccard metric is defined as follows

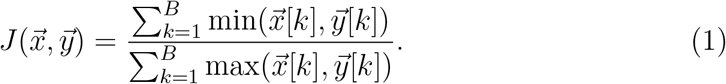

Note that by construction,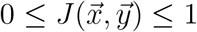. If blurry neighborhoods 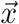 and 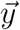 are identical, they will have Jaccard similarity of 1, and if they have 0 overlap in each bin, they will have Jaccard similarity of 0. This is a generalization of the Jaccard index of set theoretic origin, which reflects the similarity between two sets by taking the ratio of their intersection to their union.

### 2.3 Discretizing blurry neighborhood representations into a 3Dn alphabet

We can reduce the computational overhead of the blurry neighborhood-based structure comparison process by clustering neighborhoods into a 3Dn alphabet. This allows us to compare two blurry neighborhoods by looking up a score between their corresponding characters as opposed to computing the Jaccard metric. To discretize the blurry neighborhoods, we use a graph clustering approach to define 20 “landmark” vectors and associate each with a character. We then assign each amino acid to the character associated with the landmark vector that is nearest with respect to the Jaccard metric.

To construct the landmark blurry neighborhoods we cluster randomly selected 20000 blurry neighborhoods 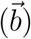 of amino acids from different protein structures. We construct a graph by connecting a pair of blurry neighborhoods 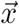 and 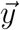 by a weighted edge with the edge weight equal to the Jaccard similarity 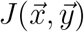 between them. To make this graph sparse, we discard the low weight edges and only keep 20 nearest neighbors for each node. Then, we use Louvain community detection algorithm with the resolution parameter tuned to give 20 final clusters that discretize the neighborhoods [2, 21]. For each cluster, we define the landmark blurry neighborhood by taking the median entry-wise of the blurry neighborhoods 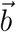 in that cluster. We can visualize the landmark blurry neighborhoods, which allows us to qualitatively view the local structure encoded by each character in the 3Dn alphabet, see Figures 3 and 4.

We considered two other approaches for discretizing our neighborhood representations: (i) training a VQVAE on the *n*-hot neighborhood vectors and (ii) further optimizing our graph cluster centers using a mutual information objective. Neither were as successful as the graph clustering approach, see Appendix A.3.

### 2.4 Constructing an alphabet based on backbone dihedral angles

As a point of comparison, we construct an alphabet derived only from the backbone dihedral angles. First, we discretize the space of *ϕ* and *ψ* backbone dihedral angles with a 30 by 30 grid, resulting in 900 bins. Many of these bins are in infeasible regions of Ramachandran plot or only account for a very small number of examples, so we reduce the number of bins to 251. Using a mutual information objective, we cluster the bins to obtain a dihedral alphabet with 20 characters. We consider the distribution over pairs of bins given by the empirical frequency of how often we observe each pair of bins substituting for each other among aligned positions in our training set of aligned structures. Our clustering algorithm iteratively merges the pair of adjacent bins that results in the distribution with maximal mutual information. See Appendix A.5 for details. Supplementary Figure 6 depicts the regions of Ramachandran plot corresponding to each character and the corresponding learned BLOSUM matrix.

### 2.5 Computing alignments

In order to determine an alignment between two proteins using the Smith-Waterman local sequence alignment algorithm [22], it is first necessary to construct a suitable similarity matrix *M*, where *M*_*ij*_ quantifies the structural similarity between the *i*^*th*^ amino acid in the first protein and the *j*^*th*^ amino acid in the second. In our blurry neighborhood method, *M*_*ij*_ is determined by a log-odds transformation of the Jaccard metric applied to the corresponding blurry neighborhoods (see Appendix A.4). For our 3Dn alphabet, we train a BLOSUM matrix on the 3Dn characters (see Appendix A.4), and let *M*_*ij*_ be the BLOSUM score for the corresponding pair of 3Dn characters. When we combine *n* different alphabets, we instead consider matrix 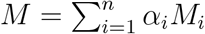,where *M*_*i*_ is the contributed matrix using alphabet *i*; the weightings *α*_*i*_ *>* 0 are determined in a search process outlined in Appendix A.2.

We construct an alignment between a pair of proteins using the Smith-Waterman algorithm [22] with the scoring matrix *M* as implemented in [19]. We used a hyperparameter search process outlined in Appendix A.2 to determine the gap open and gap extend penalties. After acquiring the local sequence alignment, we evaluate alignment quality using the Local Distance Difference Test (lDDT) [18] and use the lDDT to rank the results in our search task.

## 3 Results

### 3.1 Search benchmark

We evaluate the efficacy of our blurry neighborhood-based algorithm on a database search task using proteins from SCOPe40 (see Appendix A.6). We evaluate the lDDT of the alignment constructed between a fixed query protein and each target protein in the database, and rank the target proteins by descending lDDT value. The efficacy of each query is evaluated using sensitivity up to the first false positive across family, superfamily, and fold levels, which is the ratio of the number of true positives before the first false positive to the total number of true positives at each classification level. We average sensitivities up to the first false positive over all query proteins at each of the family, superfamily, and fold levels to benchmark our algorithm performance at different levels of divergence, see Table 2.

**Table 1:**
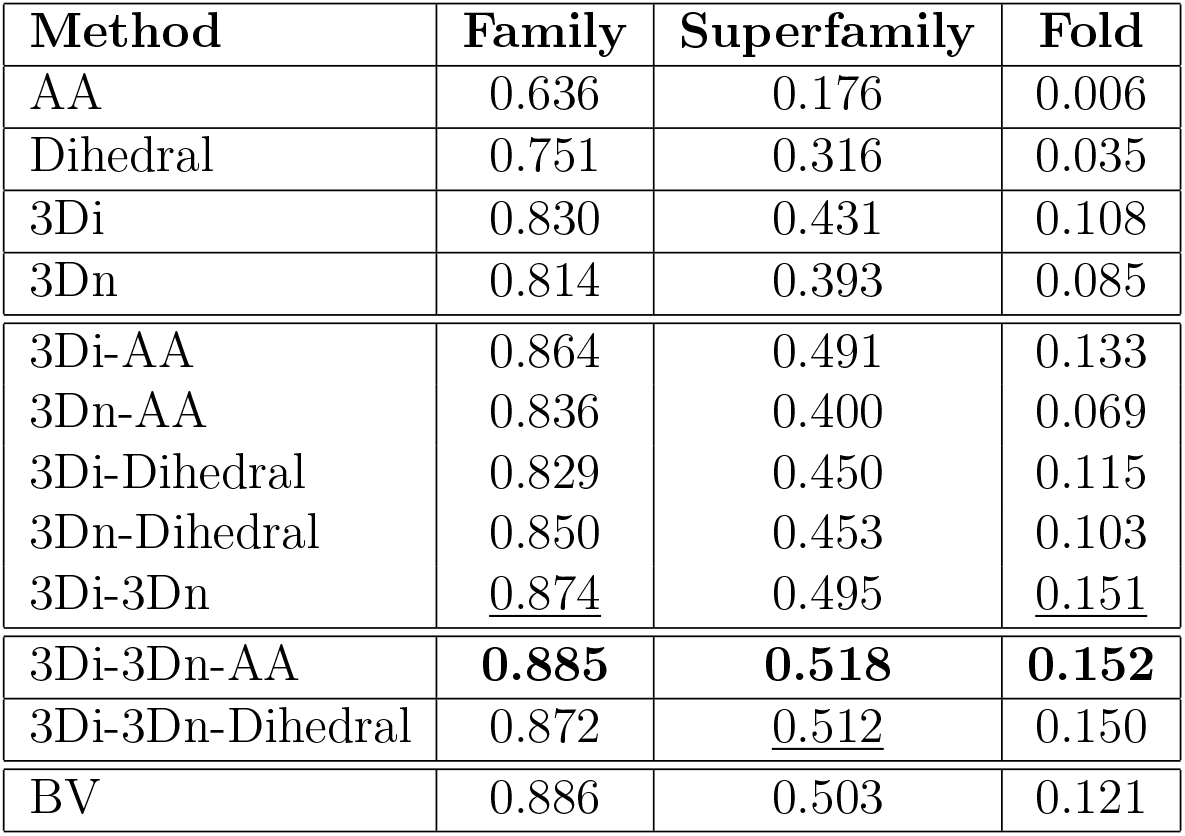
Comparison of sensitivity up to the first false positive across Family, Superfamily, and Fold categories. AA refers to the amino acid alphabet with BLOSUM62 scoring [11], Dihedral refers to an alphabet based on backbone dihedral angles described in Section 2.4, 3Di refers Foldseek’s 3Di alphabet [23], 3Dn refers our proposed alphabet, and BV refers to the scoring method using blurry neighborhood vectors with the Jaccard similarity metric. The relative weights of the combined alphabets as well as the gap penalties for each method are given in Appendix A.2. The top performing alphabet method is bolded, and the top performing alphabet method that does not require amino acid identity is underlined.

**Table 2:** Gap open, gap extend, and weights used for each method. Note that for 3Di and 3Di-AA we used the default parameters given by Foldseek.

| Method | Gap Open | Gap Extend | Weights |
| --- | --- | --- | --- |
| AA | -6.0 | -0.5 | N/A |
| Dihedral | -6.0 | -0.5 | N/A |
| 3Di* | -10.0 | -1.0 | N/A |
| 3Dn | -10.0 | -0.5 | N/A |
| BV | -8.0 | -1.0 | N/A |
| 3Di-AA* | -10.0 | -1.0 | 1.4/2.1 |
| 3Dn-AA | -6.0 | -0.5 | 0.4/0.6 |
| 3Di-Dihedral | -6.0 | -0.5 | 0.6/0.4 |
| 3Dn-Dihedral | -2.0 | -0.5 | 0.3/0.7 |
| 3Di-3Dn | -10.0 | -0.5 | 0.6/0.4 |
| 3Di-3Dn-AA | -2.0 | -0.5 | 0.4/0.2/0.4 |
| 3Di-3Dn-Dihedral | -4.0 | -0.5 | 0.4/0.3/0.3 |

Among the alphabet methods that do not require amino acid identities, the 3Di-3Dn-Dihedral and 3Di-3Dn combination alphabets are the top performers. The overall best performance is achieved by the 3Di-3Dn-AA combination alphabet. Notably, the 3Di-3Dn combination alphabet is a substantial improvement over using each alphabet individually, suggesting that the 3Di and 3Dn alphabets encode different information. We explore these differences in the following section. Moreover, the performance of the 3Di-3Dn and the 3Di-3Dn-Dihedral alphabets are modest improvements over the 3Di-AA alphabet combination used by Foldseek. Thus, we have established a method that does not use amino acid identities but gives comparable results to Foldseek, which is of interest for structure search when homology is not expected.

We also consider the performance of the Blurry Vector (BV) approach, which is an instantiation of our blurry neighborhood method with the discretization utilizing 1000 bins. Our BV method performs similarly to the 3Di-3Dn combination alphabet while also not requiring amino acid identities. However, our BV method is much less efficient than the alphabet methods because the similarity matrix is computed with a Jaccard metric rather than computed via look-ups in a BLOSUM matrix. We include the BV results to draw attention to the discrepancy in performance between the BV method and 3Dn alphabet, suggesting there is a substantial loss in information in the clustering process.

Figures 2A and 2B illustrate the performance gains achieved by augmenting the 3Di, 3Dn, and combined 3Di-3Dn alphabets with the amino acid and dihedral alphabets, as measured by the average sensitivity up to the first false positive for the Superfamily search. Figure 2C gives the precision recall curves at the superfamily level. Analogous figures for the Family and Fold levels are given in Supplementary Figures 7 and Figure 8.

**Figure 1:**
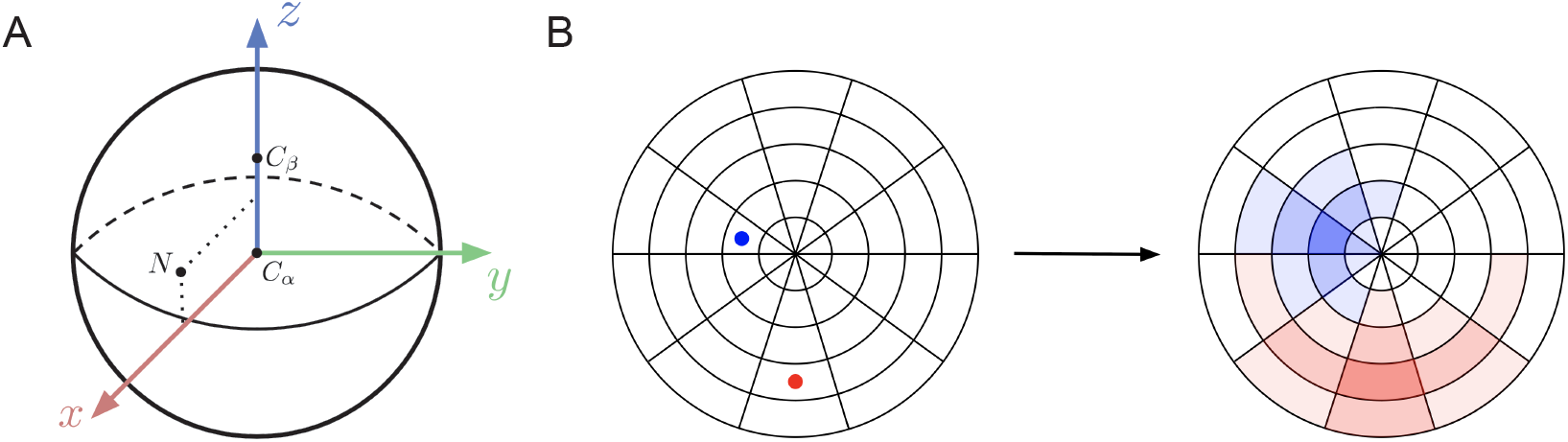
Constructing blurry neighborhoods. A. Visualization of the reference frame generated during the *n*-hot vector construction process, with the central amino acid at the origin. B. Effect of applying the transition matrix to the *n*-hot vector, showing only the bins that cross the *x*-*y* plane. The left image depicts an *n*-hot vector representing a neighborhood with one beta sheet neighbor (blue dot) and one right helix neighbor (red dot). The right image represents the blurry neighborhood obtained applying the transition matrix to the *n*-hot vector. Stronger blue or red color presence in a bin represents a higher probability that the respective amino acid migrates to that bin in one evolutionary timestep.

**Figure 2:**
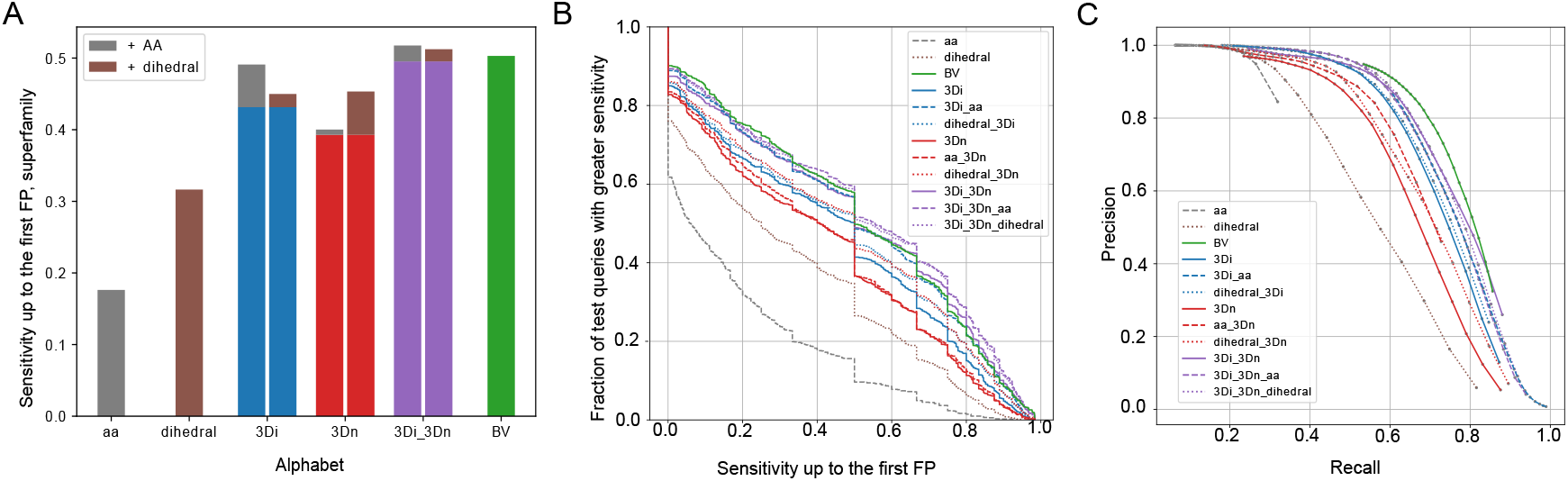
Superfamily search benchmark results. A. Sensitivity up to the first false positive for AA, Dihedral, 3Di, 3Dn, 3Di-3Dn, and BV alphabets at the superfamily level. The benefit of adding amino acid information is shown in gray, and the benefit of adding dihedral information is shown in brown. B. The fraction of queries that receive a greater sensitivity up to the first false positive than indicated in the corresponding *x*-axis value, at the superfamily level. Each color and line type represent a different alphabet or combination of alphabets. C. Precision and recall of each alphabet at the superfamily level. Each color and line type represent a different alphabet or combination of alphabets.

**Figure 3:**
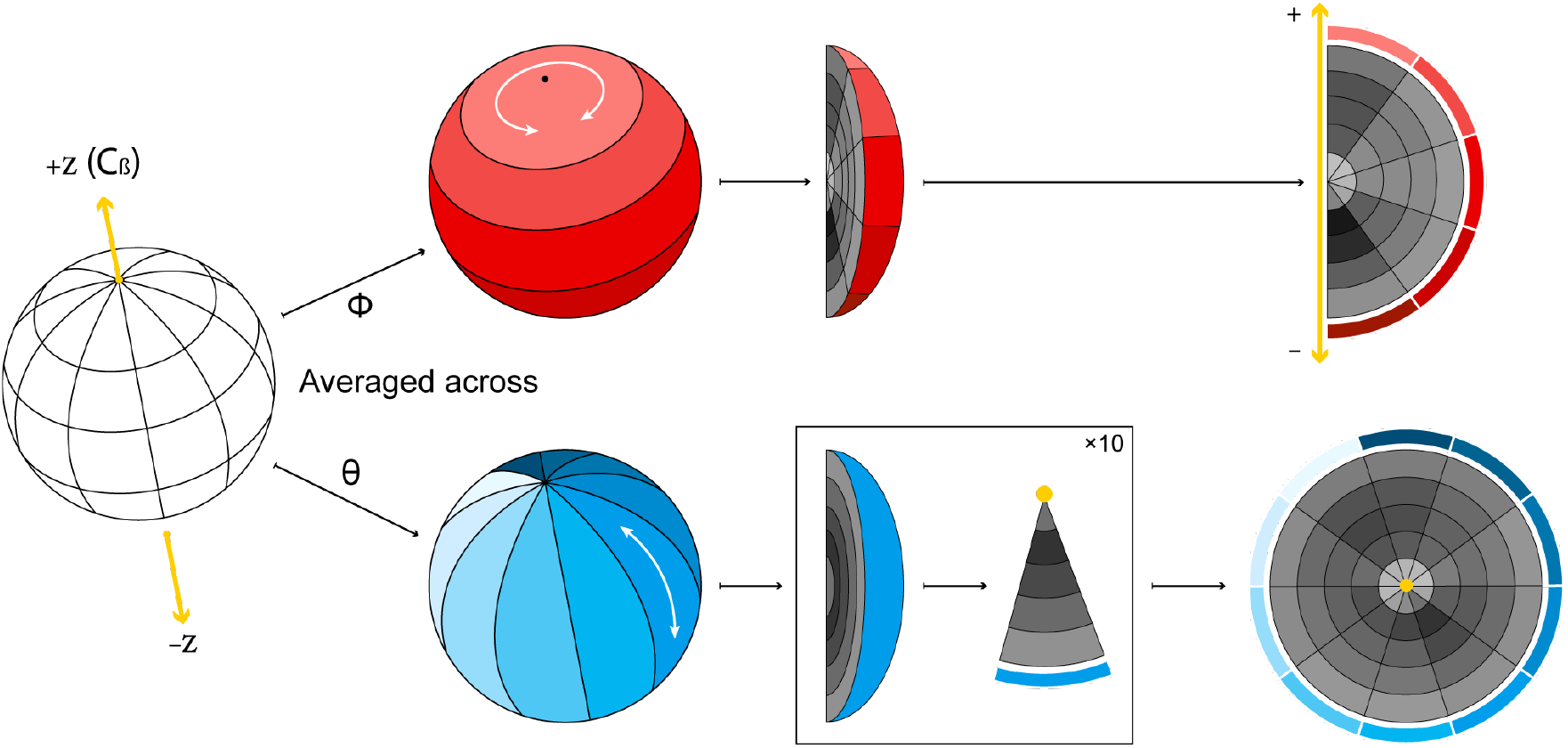
Visualizing 3Dn characters. We discretize the 15Å sphere oriented with respect to the reference frame depicted in Figure 1 into 250 bins by taking 10 intervals in *ϕ*, 5 intervals in *θ*, and 5 intervals in *r* (hidden in the left image). To visualize the neighbor counts in each of these bins, we show two flattened images obtained by averaging over *ϕ* (upper row) and *θ* (lower row). In the upper row, the color of each cell represents the average number of neighbors across all bins with a particular interval for *r* and interval for *θ*. In the lower row, the color of each cell represents the average number of neighbors across all bins with a particular interval for *r* and interval for *ϕ*. The *z*-axis appears as a line in the upper plot and a point in the lower plot, as depicted in yellow. See Figure 4 for examples. Note that these figures are not to scale; true *r, θ*, and *ϕ* values are chosen so that each bin has equal volume.

**Figure 4:**
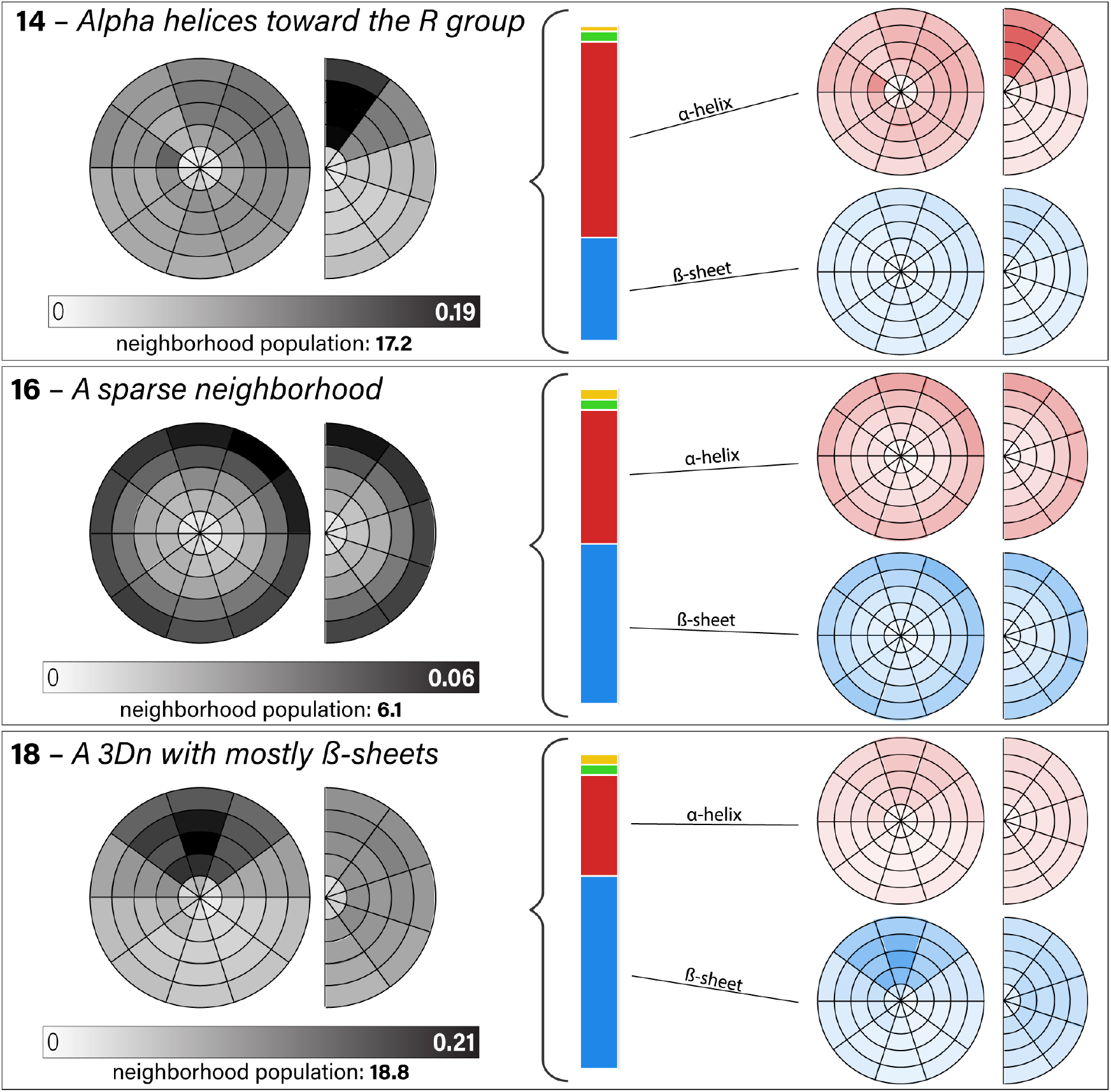
Visualization of 3Dn characters 14, 16, and 18. Figure 3 explains how to interpret the circle and semi-circle figures. The left gray figures illustrate the spatial distributions of neighbors in the landmark blurry neighborhood for the 3Dn character. The number of neighbors varies substantially across the different 3Dn states, as indicated by the given neighborhood populations. The vertical bar shows the distribution of the secondary structure of the neighbors. The top rightmost plots illustrate the distribution of the right helix neighbors, whereas the bottom rightmost plots illustrate the distribution of beta sheet neighbors. See Section 3.2.1 for a qualitative description of the three 3Dn states visualized here. The full alphabet, including left *α*-helix (yellow) and unclassified secondary structure (green) is depicted in Figure 9.

### 3.2 Interpretation of the 3Dn alphabet

#### 3.2.1 Visualization of 3Dn characters

We can visualize each 3Dn character with two figures, constructed by averaging over bins delimited by *ϕ* (creating a semi-circle) and by *θ* (creating a circle) as shown in 3. Figure 4 depicts three 3Dn characters: 14, 16, and 18. The 3Dn character 14 is representative of many characters in our alphabet, detecting a particular secondary structure (in this case, right *α*-helices) toward (or away from) the *R* group. The 3Dn character 16 represents the sparsest neighborhoods with 6.1 neighbors total. The 3Dn character 18 shows *β*-sheet neighbors uniformly distributed with respect to the angle to the *R* group, but with a distinct pattern with respect to the orientation in our *XY* plane.

#### 3.2.2 Comparison of alphabets: AA, 3Di, Dihedral, 3Dn

In order to better understand the biochemical features encoded by Foldseek’s 3Di alphabet and our 3Dn alphabet, we analyze the co-occurrance of amino acid identity, dihedral cluster, and 3Dn and 3Di characters. We find that 3Dn characters encode amino acid identity more than 3Di characters, and 3Di characters encode dihedral information more than 3Dn characters. This finding is consistent with the observation that the 3Dn alphabet exhibits greater improvement when augmented with the dihedral alphabet as compared with the amino acid alphabet, and vice versa for the 3Di alphabet (see Figure 2A).

First, we explore the relationship between 3Dn and 3Di characters. Recall from Table 2 that combining 3Dn and 3Di alphabets on the search benchmarking tasks outperforms the individual performance of the 3Dn and 3Di alphabets, suggesting that 3Dn and 3Di alphabets capture different structural information. We confirm this intuition by analyzing the co-occurrence of 3Dn and 3Di characters. We compute the ratio between how often each 3Di and 3Dn character pair co-occurs (e.g. how many times we observe an amino acid assigned to the particular pair of 3Di and 3Dn characters) and how often we would expect for them to co-occur under the null distribution that 3Di and 3Dn characters are assigned at random with probability given by their empirical frequencies. Figure 5A, depicts the log of these ratios; the dark blue regions indicate pairs that rarely co-occur, and the red indicates pairs that co-occur more than would be expected by chance. There is no clear mapping between 3Di and 3Dn characters, suggesting that the characters capture different structural information.

**Figure 5:**
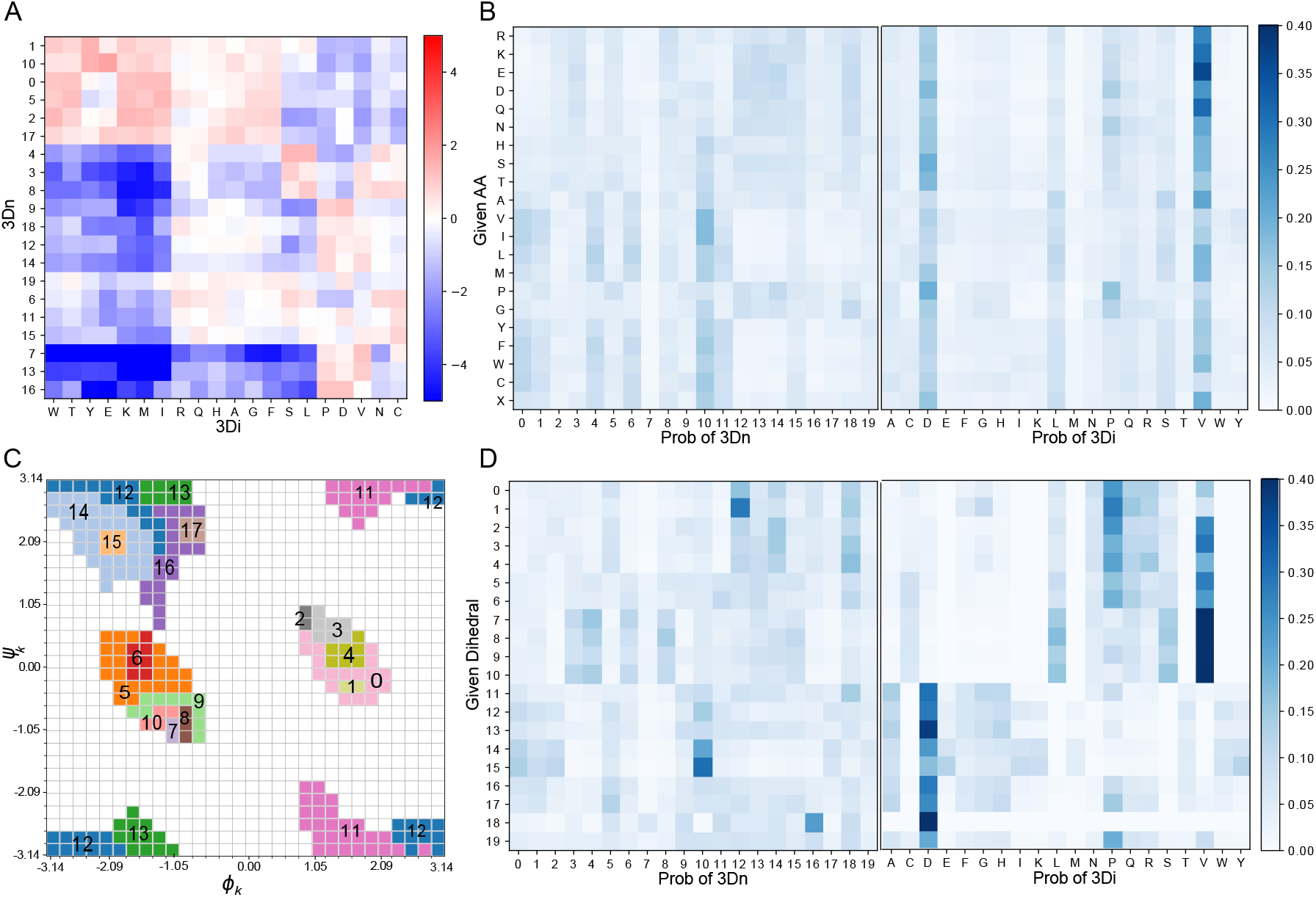
Co-occurance of 3Di, 3Dn, amino acid and dihedral characters. A. A log odds matrix illustrating the co-occurance of 3Di and 3Dn characters; a positive log odds score (red) indicates the pair co-occur more than what would be expected by chance. B. Probabilities of any given amino acid belonging to any 3Dn character or 3Di character. C. Labels of regions corresponding to our learned dihedral characters on a Ramachandran plot. D. Probabilities of any given dihedral character associating to any given 3Dn character or 3Di character.

**Figure 6:**
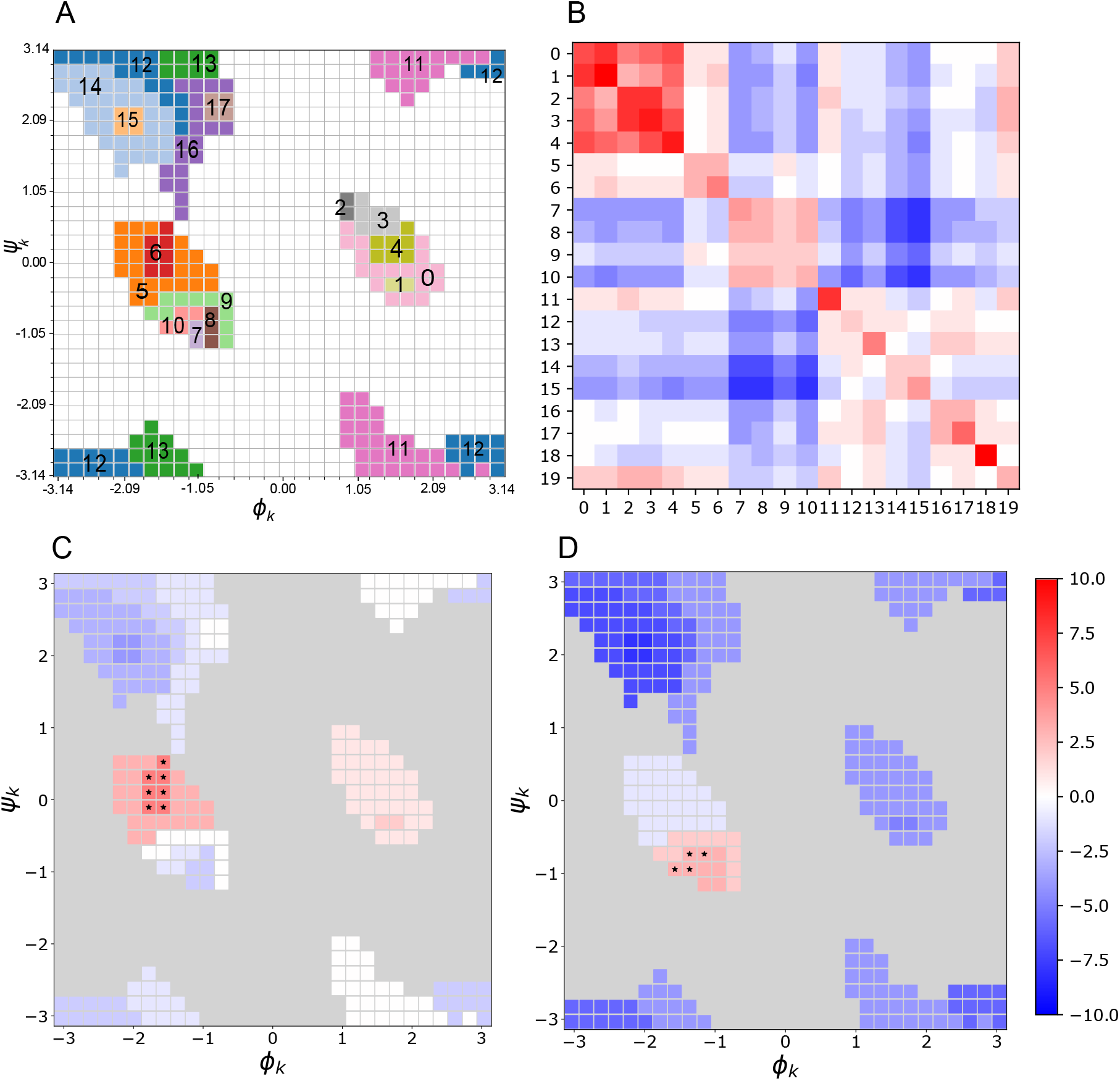
Backbone dihedral alphabet and corresponding BLOSUM matrix. A. Labeled clusters on a Ramachandran plot. B. BLOSUM-like substitution matrix for dihedral alphabet. C and D. Substitution scores between a chosen cluster *i* (indicated by a set of stars) and all other clusters visualized on a Ramachandran plot, with *i* = 6 in C and *i* = 10 in D.

**Figure 7:**
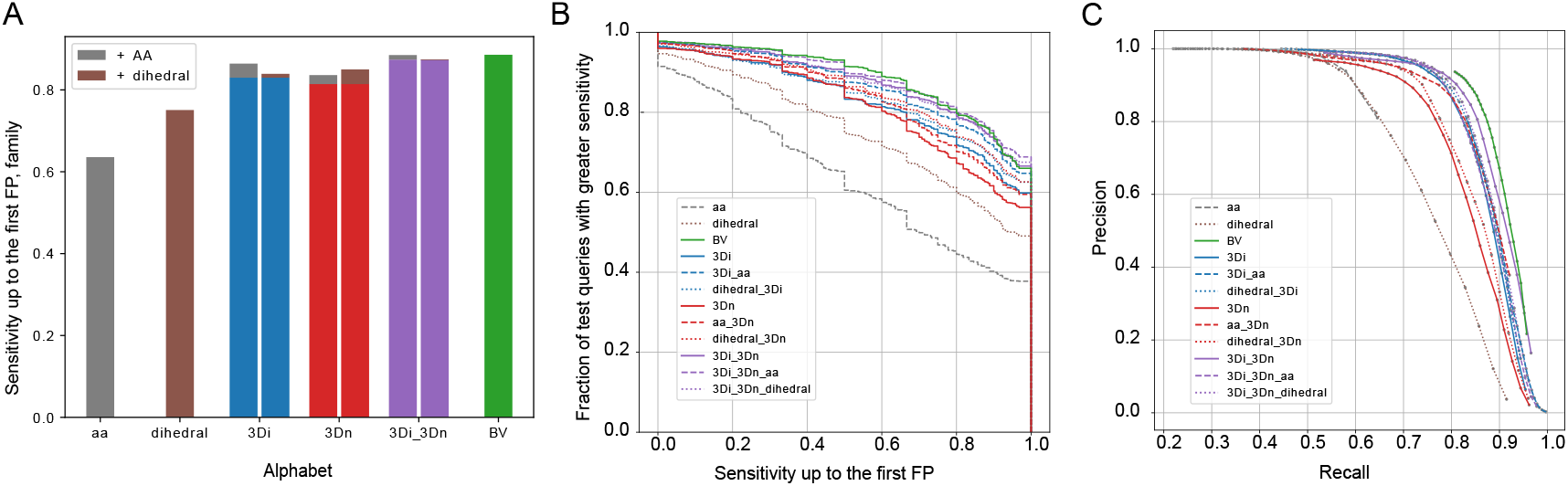
Family search benchmark results. A. Sensitivity up to the first false positive for AA, Dihedral, 3Di, 3Dn, 3Di-3Dn, and BV alphabets at the family level. The benefit of adding amino acid information is shown in gray, and the benefit of adding dihedral information is shown in brown. B. The fraction of queries that receive a greater sensitivity up to the first false positive than indicated in the corresponding *x*-axis value, at the family level. Each color and line type represent a different alphabet or combination of alphabets. C. Precision and recall of each alphabet at the family level. Each color and line type represent a different alphabet or combination of alphabets.

**Figure 8:**
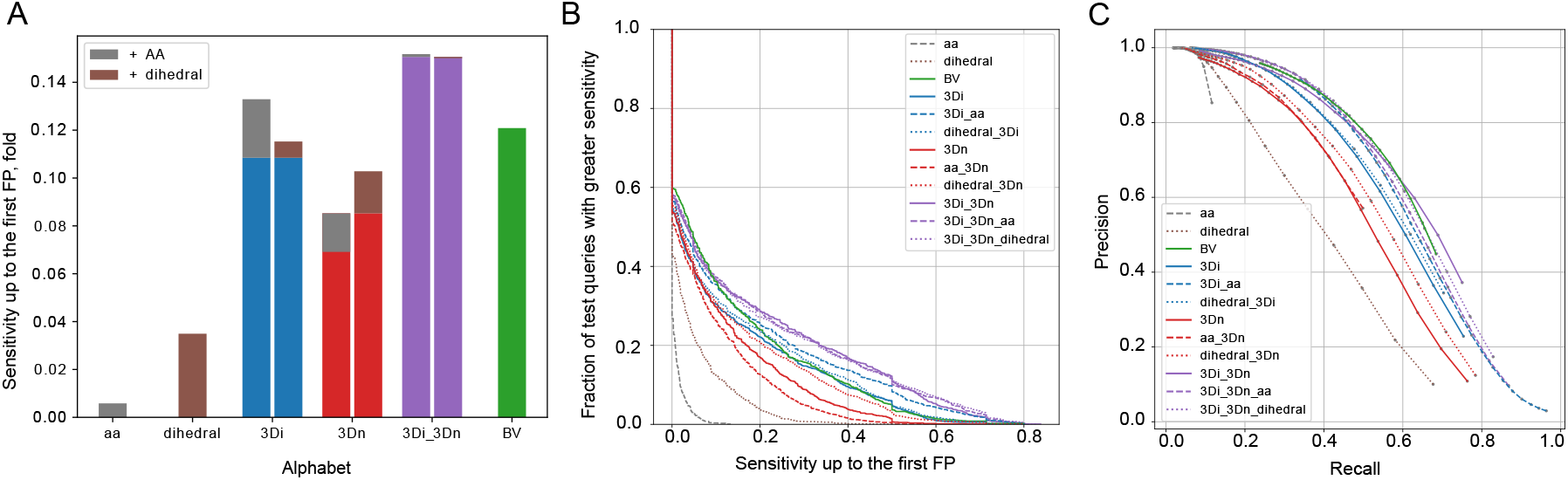
Fold search benchmark results. A. Sensitivity up to the first false positive for AA, Dihedral, 3Di, 3Dn, 3Di-3Dn, and BV alphabets at the fold level. The benefit of adding amino acid information is shown in gray, and the benefit of adding dihedral information is shown in brown. B. The fraction of queries that receive a greater sensitivity up to the first false positive than indicated in the corresponding *x*-axis value, at the fold level. Each color and line type represent a different alphabet or combination of alphabets. C. Precision and recall of each alphabet at the fold level. Each color and line type represent a different alphabet or combination of alphabets.

Next we compare the co-occurrence of 3Dn and 3Di characters to amino acid identity to evaluate the extent to which each alphabet encodes amino acid identity (Figure 5B). We observe that certain 3Dn characters tend to exclusively represent polar amino acids or exclusively represent nonpolar amino acids. For instance, 3Dn characters 12, 13, and 14 are more common among polar amino acids, and 3Dn characters 4 and 6 are more common among nonpolar amino acids. Such patterns are not as prominent among the 3Di alphabet, perhaps explaining why when amino acid information is added, 3Di search performance is substantially boosted whereas 3Dn search performance only marginally improves (see gray bars in Figure 2A).

Similarly, Figure 5D illustrates the extent to which the 3Di and 3Dn alphabets encode secondary structure, as measured by backbone dihedral angles. Figure 5C depicts our dihedral alphabet; the discretization and subsequent clustering process based on mutual information is detailed in Section 2.4 and Appendix A.5. We observe that the backbone dihedral angles and consequently, secondary structure is reflected in 3Dn characters as well, with 3Dn character 10 common among beta sheets, characters 3 and 4 common among right helices, and character 12 common among left helices. Recall from Figure 2A that including dihedral information provides a larger performance enhancement for 3Dn characters than 3Di characters in the search benchmark. This may be because 3Di characters appear to already capture more dihedral information than 3Dn characters, indicated by stronger colored regions in the corresponding graph in Figure 5D. Therefore, including dihedral information with 3Di characters may be redundant in terms of information captured, accounting for the smaller improvement in combining 3Di and dihedral alphabets when compared to combining dihedral and 3Dn alphabets.

Interestingly, the matrices in Figure 5D differentiate regions in the Ramachandran plot in a way that agrees with the patterns of conservation encoded in the learned dihedral BLOSUM matrix. Although dihedral characters 5, 6, 7, 8, 9, and 10 all correspond to adjacent regions on the Ramachandran plot we witness that characters 5 and 6 have strikingly different relationships to the 3Dn characters than characters 7, 8, 9, and 10, with a similar yet weaker trend also holding for 3Di characters. This suggests that amino acids with dihedral angles in region 5 and 6 form different structural relationships than those in 7-10. This is supported by the dihedral BLOSUM matrix which illustrates that the substitution pattern for characters 5 and 6 differ from that of 7-10, see Supplementary Figure 6.

#### 3.2.3 Comparison of performance by protein characteristics

To further gain insight on the limitations of 3Di and 3Dn alphabets on structural comparison tasks, we investigated whether they perform differently depending on the characteristics of the proteins being aligned. In Figure 10, we consider the performance of 3Di and 3Dn methods on query proteins depending on their protein class, a level of protein classification weaker than fold. Here, we find that at family, superfamily, and fold levels, 3Dn performs better with smaller proteins, and at the family level, the 3Dn alphabet performs better with alpha helix dominant proteins. Alternatively, we see better performance at the family and superfamily level by the 3Di alphabet on beta sheet dominant proteins. While further investigation is necessary to fully explain these discrepancies we offer some hypotheses. The 3Dn alphabet encodes structural information on a larger scale spatially than 3Di; 3Dn characters capture all amino acids that are more than 5 positions away in the sequence and with 15Å. Since alpha helices are more dense than beta sheets, comparatively more helix information may be encoded in the neighborhood. For small proteins, many amino acids are on the boundary of the protein and will have 15Å neighborhoods with large unoccupied regions, which can be expressed by 3Dn characters.

**Figure 9:**
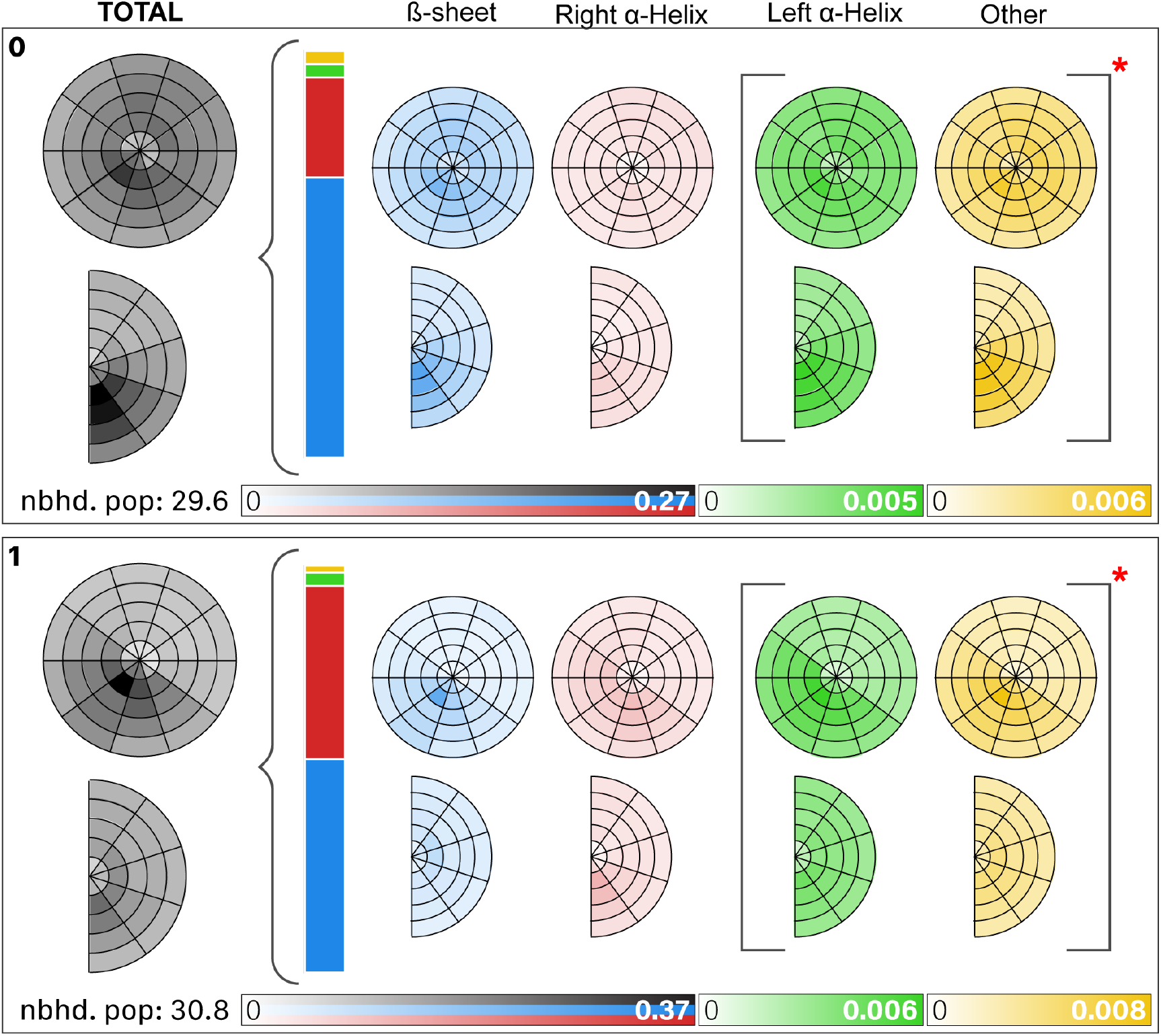
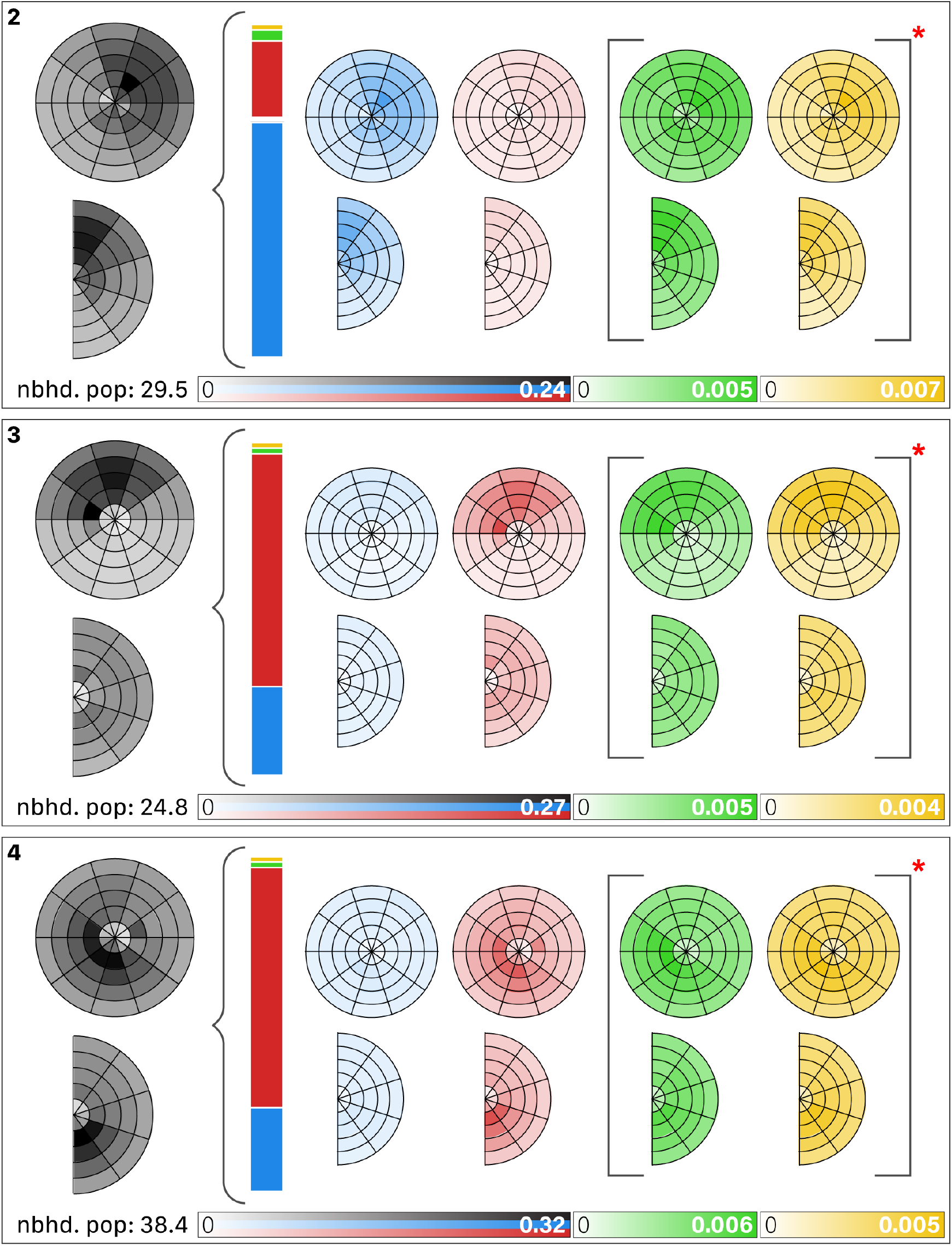
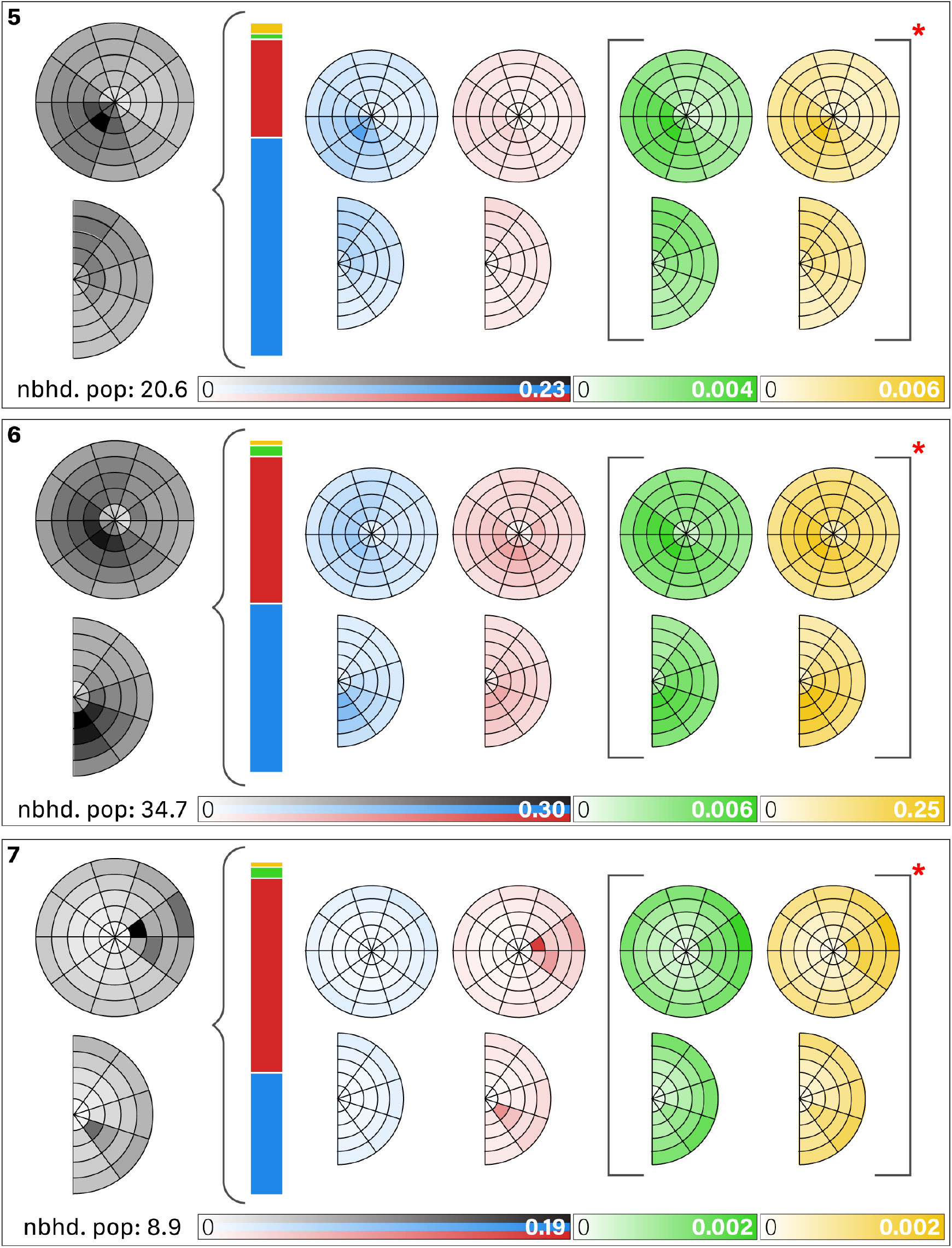
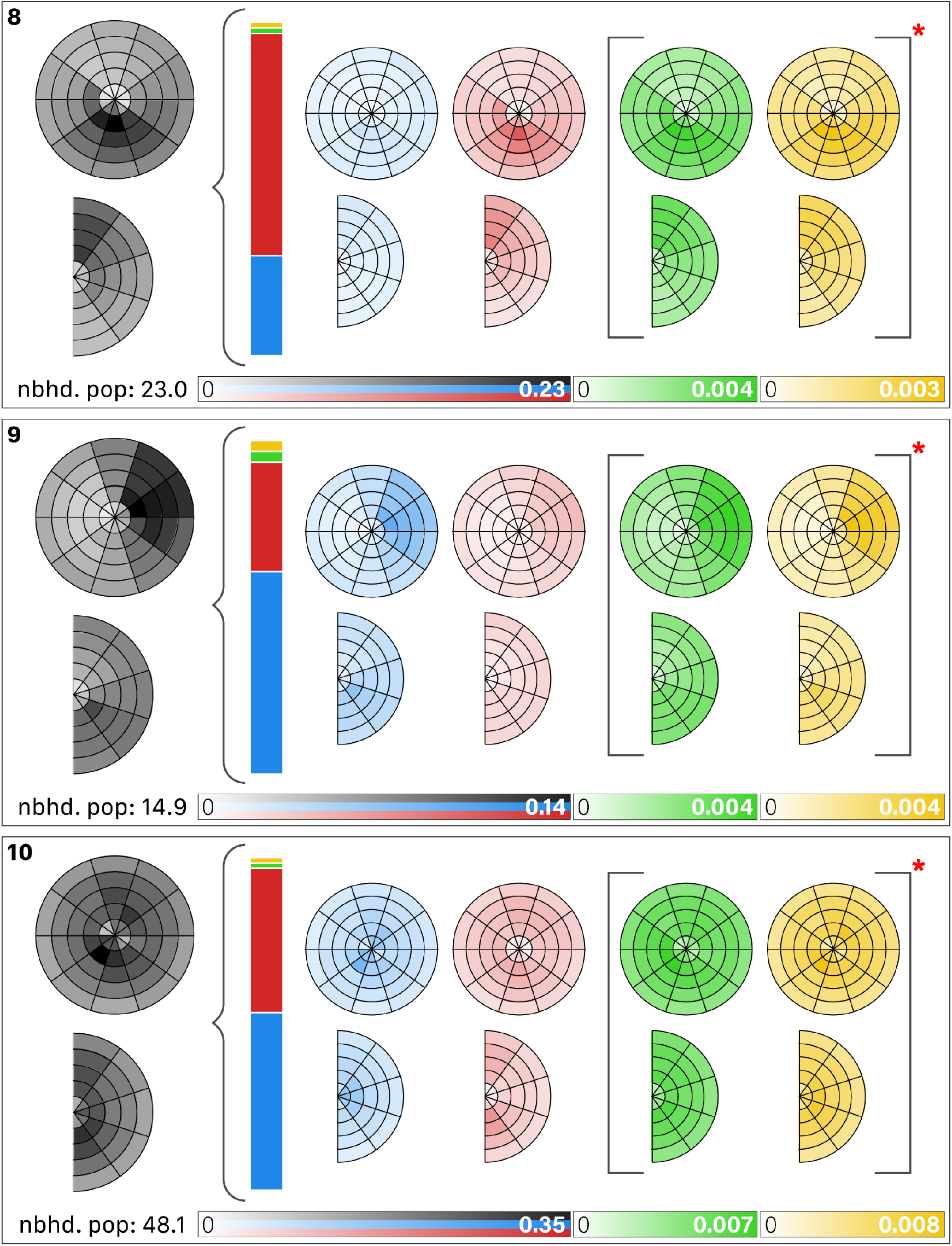
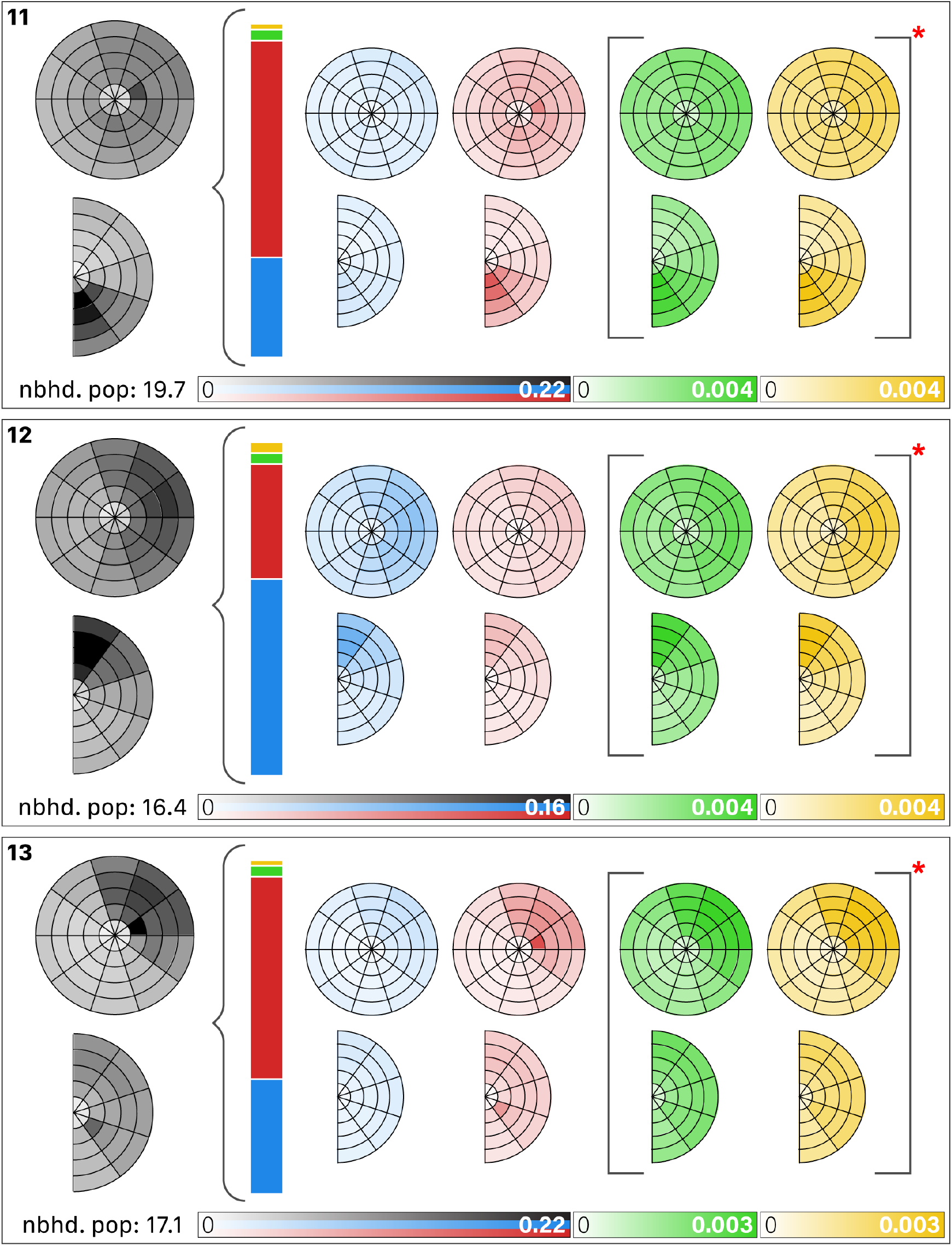
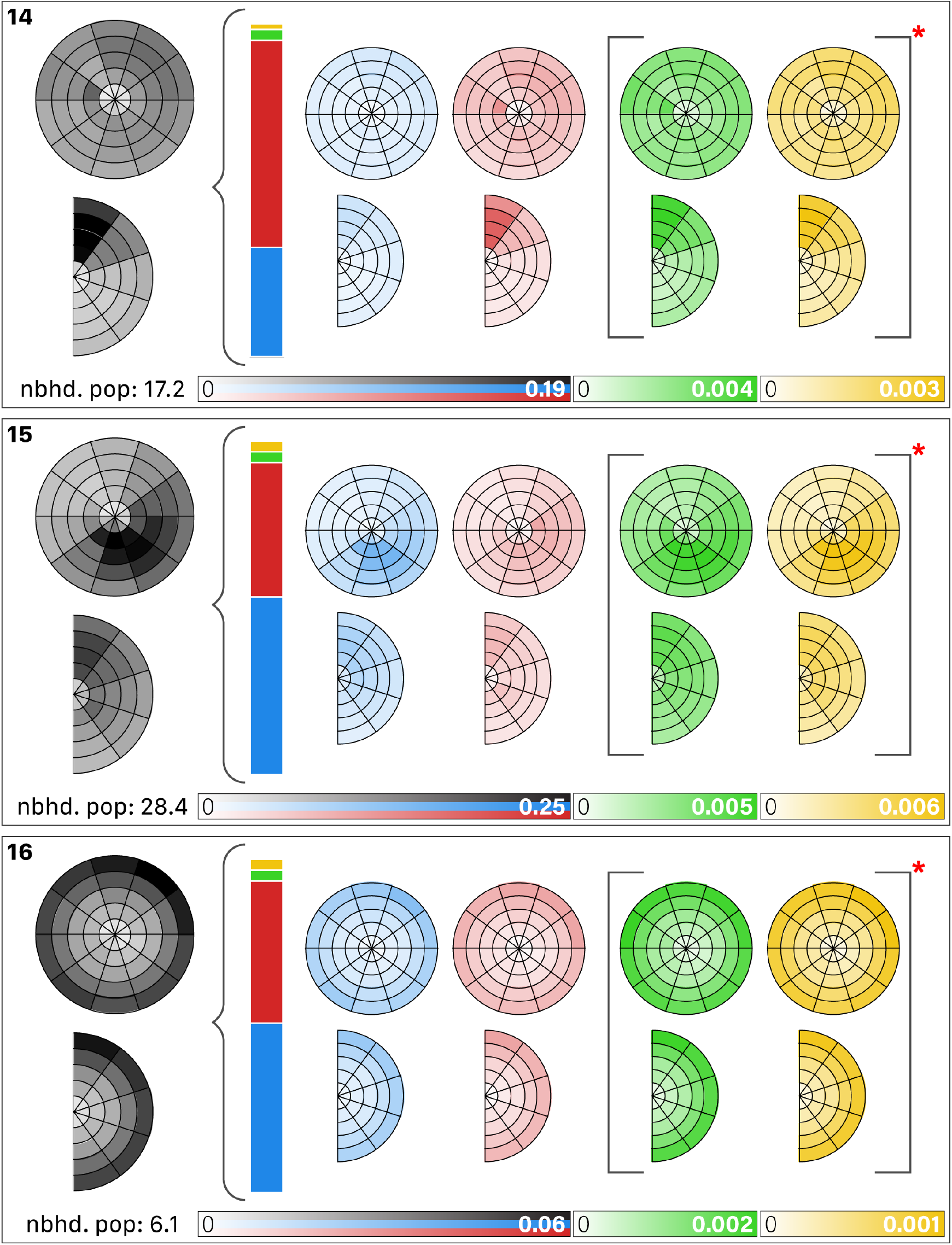
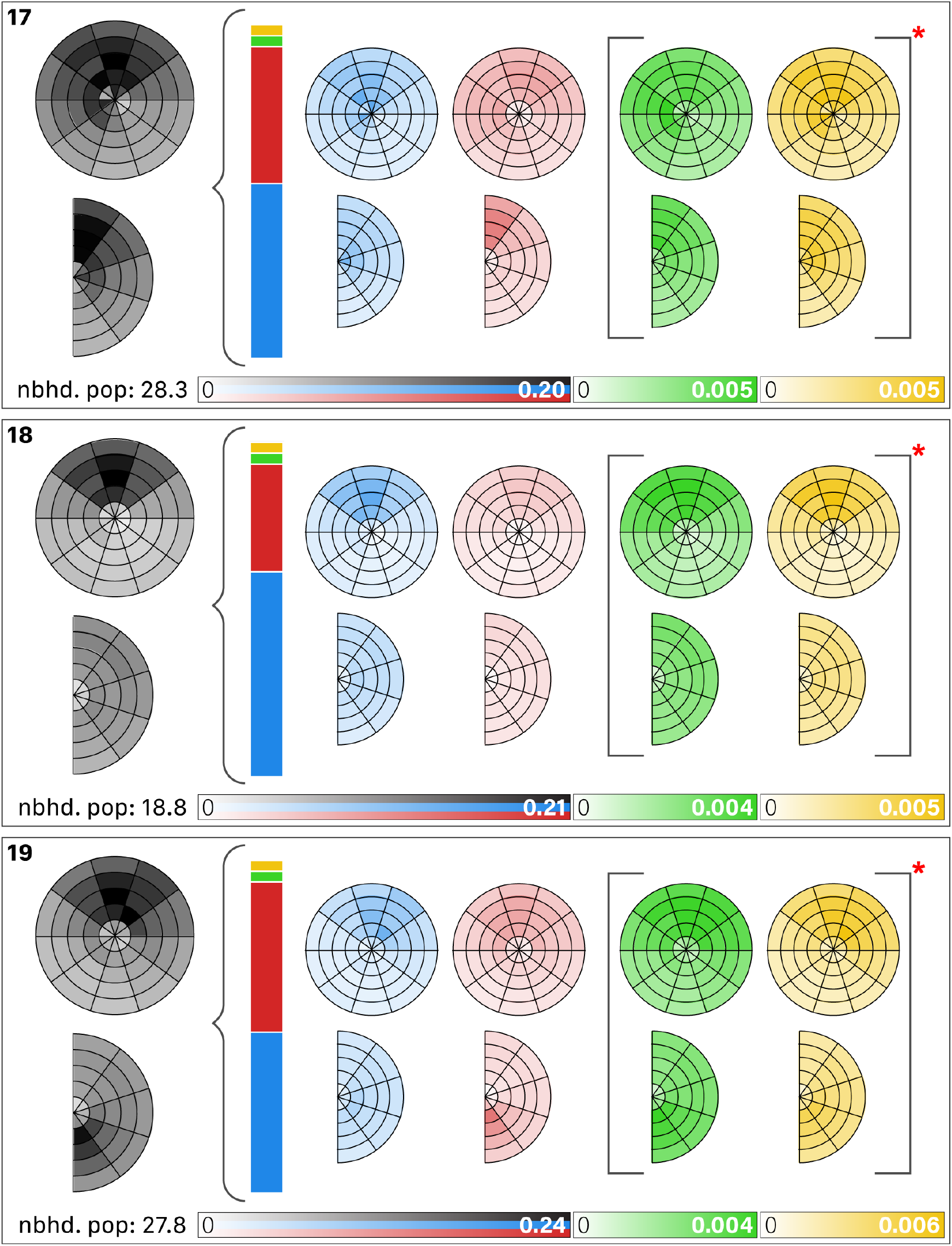
The complete 3Dn alphabet. Figure 3 explains how to interpret the circle and semi-circle figures. The left gray figures illustrate the spatial distributions of neighbors in the landmark blurry neighborhood for the 3Dn character. The number of neighbors varies substantially across the different 3Dn states, as indicated by the given neighborhood populations and different scales of the black, blue, and red colorbar between different 3Dn characters. The four pairs of circles and semicircles depict the distribution of neighbors by secondary structure type; from left to right: beta sheet (blue), right alpha helix (red), left helix (green), unclassified (yellow). We use a different colorbar scale (normalized to the highest value) to visualize left *α*-helix and unclassified to make the patterns visible.

**Figure 10:**
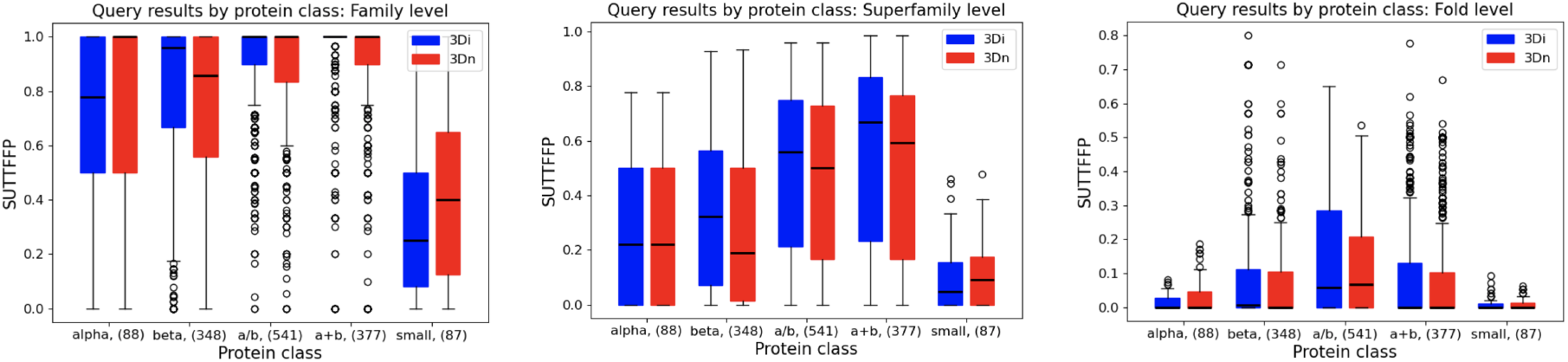
Query results by protein class at family, superfamily, and fold levels in the search benchmarking task. Query results are quantified by sensitivity up to the first false positive (SUTTFFP). Protein class refers to the class of the SCOPe ID of the query protein: ‘alpha’ refers to proteins dominated by alpha helices, ‘beta’ refers to proteins dominated by beta sheets, ‘a/b’ refers to proteins where alpha helix and beta sheet components exist in separate, continuous regions in the protein, ‘a+b’ refers to proteins that contain alternating alpha helix and beta sheet components within the protein, and ‘small’ refers to small proteins. The number of queries in the class is noted in parentheses. In the box plot, the bottom edge of the box is at *x*_25_, the upper edge is at *x*_75_, the middle bar represents the median value *x*_50_, the bottom whisker is at max(0, *x*_25_ *−* 1.5(*x*_75_ *− x*_25_)), and the top whisker is at min(1.0, *x*_75_ + 1.5(*x*_75_ *− x*_25_)), where *x*_*p*_ represents the value at percentile *p*.

We also consider the performance of 3Di and 3Dn alphabets by considering the alignment quality of pairs. Figure 11 shows the distribution of lDDTs for pairs of various protein classes. We find again that 3Dn performs better on smaller proteins. Additionally, we considered alignment quality by relative protein length, which we defined as 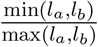, where *l*_*a*_ and *l*_*b*_ refer to the sequence lengths of proteins *a* and *b*, respectively. Here, we do not see any clear performance trends between 3Di and

**Figure 11:**
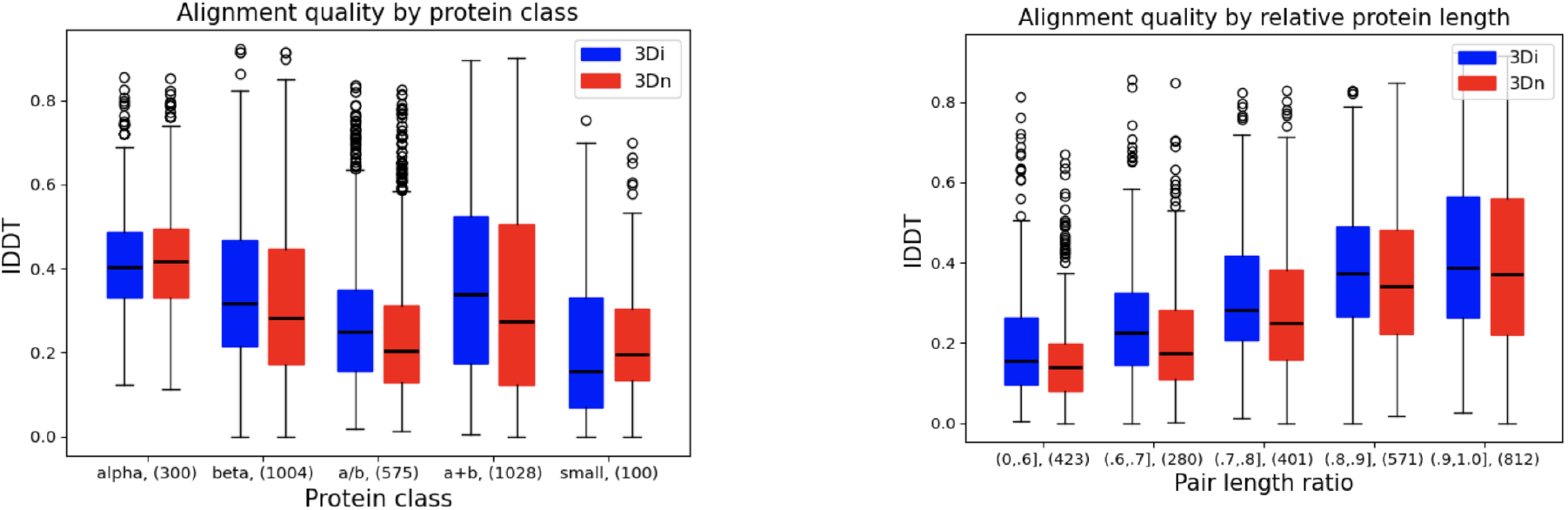
Alignment quality on the test set for 3Di vs. 3Dn with respect to various protein qualities. The figure on the left depicts lDDT of protein pairs with respect to the class of the proteins within the pair, according to their SCOPe IDs: ‘alpha’ refers to proteins dominated by alpha helices, ‘beta’ refers to proteins dominated by beta sheets, ‘a/b’ refers to proteins where alpha helix and beta sheet components exist in separate, continuous regions in the protein, ‘a+b’ refers to proteins that contain alternating alpha helix and beta sheet components within the protein, and ‘small’ refers to small proteins. The number of pairs in the class is noted in parentheses. The figure on the right depicts lDDT with respect to the ratio of the sequence length of the shorter protein in the pair to the sequence length of the longer protein in the pair. Each bar in the plot represents lDDT of pairs which length ratio lies in the corresponding interval indicated on the x-axis, and the number of pairs in the pair length classification is noted in parentheses. In the box plot, the bottom edge of the box is at *x*_25_, the upper edge is at *x*_75_, the middle bar represents the median value *x*_50_, the bottom whisker is at max(0, *x*_25_ *−* 1.5(*x*_75_ *− x*_25_)), and the top whisker is at min(1.0, *x*_75_ + 1.5(*x*_75_ *− x*_25_)), where *x*_*p*_ represents the value at percentile *p*.

3Dn alphabets on pair alignments as a function of the relative protein lengths.

## 4 Discussion

We proposed an interpretable method for characterizing local structure in proteins by considering the spatial distributions of nonadjacent amino acid neighbors. Our blurry neighborhood method and corresponding 3Dn alphabet performed well on protein database search tasks, and when combined with 3Di characters from Foldseek, outperformed both the 3Di and the 3Dn alphabets individually. The combined 3Dn-3Di alphabet is the state of the art alphabet for local search that does not rely on amino acid identity.

Although we have demonstrated that the combined 3Dn-3Di alphabet is high performing, pursuing formal integration with search software such as Foldseek introduces a series of engineering challenges. First, a limitation of our 3Dn alphabet is that computing the 3Dn characters requires constructing a 1000-dimensional blurry vector. A 3Dn alphabet dervived from a coarser discretization of the sphere has the potential to be comparably expressive while computationally more efficient. Second, the search algorithm performance of the combined alphabets may benefit from alternative ways of combining the scores, e.g. taking a geometric mean. Third, further investigation into the ranking criteria is merited. Here we rank by lDDT, but Foldseek ranks by a combination of lDDT, TM-score, and bitscore. Fourth, when considering a protein query for a database search, Foldseek involves prefiltering steps to reduce the size of the target set, which is not currently a feature of our work. It remains to be explored whether a combination alphabet improves prefiltering or whether either individual alphabet will suffice.

More broadly, we hope that our work fuels the exploration of novel alphabets and combinations of alphabets for protein structure search. The demonstration that our 3Dn-3Di alphabet outperforms the 3Di alphabet alone establishes proof of principle that protein structure comparison can be improved with more expressive alphabets. Of particular interest is whether recent methods of tokenizing protein structures designed for deep learning [17, 8, 7] could be repurposed for protein structure comparison. We have developed a series of software tools that allow for easy construction of BLOSUM matrices for arbitrary alphabets, a mutual information-based clustering algorithm to reduce size of alphabets, and a benchmark for assessing performance on a search task, all of which can be applied to formulate and test other alphabets.

## Acknowledgements

We thank Sean Eddy and his lab where this project began. We thank Sergey Ovchinnikov, Martin Steinegger, Johanees Söding, Jeanne Trinquier, Michel van Kempen, and Sukhwan Park for useful conversations and feedback. We thank Yu-Shan Lin and Tiffani Hui for useful conversations about Ramachandran plots.

This work was supported by the NSF-Simons Center for Mathematical and Statistical Analysis of Biology at Harvard (award number #1764269) and a Burroughs-Wellcome Careers at the Scientific Interface (CASI) award. The authors acknowledge the Tufts University High Performance Compute Cluster (https://it.tufts.edu/high-performance-computing) which was utilized for the research reported in this paper. Additionally the authors acknowledge the compute resources provided by the Data Intensive Studies Center at Tufts.

## A Extended Methods

### A.1 Neighborhood construction

We encode spatial location of the neighbors of *x* by discretizing the 15Å sphere oriented with respect to the reference frame centered at the alpha carbon of *x*. We create “bins” of equal volume characterized by ranges of *R, φ*, and *θ* values, the parameters of the spherical coordinate system. We partition [0, *R*_MAX_], [0, 2π], and [0, *π*] into *N*_*R*_, *N*_*φ*_, and *N*_*θ*_ subintervals. We define the *i*^th^ subinterval, 0 indexed, of [0, *R*_MAX_] as 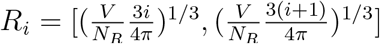,where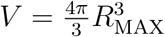. We define the *i*^th^ interval of [0, *π*] as 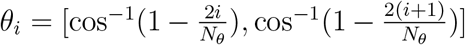. We define the *i*^th^ interval of *φ* as 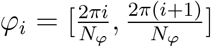.

To encode secondary structure assignment, we utilize set *S* = *{β, α*_*R*_, *α*_*L*_, *X}* to represent beta sheets, right alpha helices, left alpha helices, and an unknown secondary structure *X*, respectively. We assign an element of the secondary structure based on the *φ* and *ψ* angles of the corresponding neighbor, with *β* assigned if *φ ∈* [*−*3.14, 0.42], *ψ ∈* [0.63, 3.14] or *φ ∈* [ *−*3.14, 0.84], *ψ ∈* [ *−*3.14, 2.09], *α*_*R*_ assigned if *φ ∈* [ *−*2.51, 0.42], *ψ ∈* [ *−*1.47, 0.63], *α*_*L*_ assigned if *φ ∈* [*−*0.63, 2.51], *ψ∈* [ *−*0.84, 1.26], and *X* assigned otherwise. Our assignments are imperfect since backbone dihedral angles are not a perfect proxy for secondary structure. For instance, amino acids in turns may be assigned to helices, and the first and last amino acid will always be assigned to the extraneous *X* bin.

In our blurry neighborhood vectors, each position or “bin” corresponds to the Cartesian product of the three intervals in conjunction with the secondary structure set: *R*_*i*_ *× φ*_*j*_ *×θ*_*k*_ *× S*. Once an amino acid *y* is determined to be a neighbor of reference amino acid *x, y* will be assigned a corresponding bin. This bin reflects both the location of *y* relative to *x* and the secondary structure information of *y*.

### A.2 Validation task to choose weightings and gap penalties

We followed a two step procedure to find the best gap open and extend penalties for each alphabet. To do so, we first divided our validation set into two parts. We identified 93 proteins in the validation set that satisfy the hierarchical condition for search (the data contains at least one member in the same superfamily that is not in the family and at least one member in the same fold that is not in the same superfamily). We call this set the “validation query set.” To arrive at the “validation alignment set” we considered all TM-aligned pairs of validation proteins in which both proteins had length less than 512 and neither protein is part of the validation query set. We determined the optimal gap-open and extend as follows:

1. We align all pairs in the validation alignment set with all combinations of gap open parameters *{−*20, *−*18, *… −*2, 0*}* and gap extend parameters *{−*3, *−*2.5, *−*2, … 0*}*. We compute the gap open and extend parameters that give us the best mean lDDT of the resulting alignments and the best spearman correlation between the lDDT of our alignments and the lDDT of the TM-alignments. These parameters tended to differ from each other, and it is not clear apriori which is a better metric to use. For the validation search task, we considered all pairs of open and extend parameters that fell between these two different notions of optimal open and extend (e.g. if the optimal via mean was (*a, b*) and the optimal via spearman correlation was (*c, d*) with *a < c* and *b < d*, we considered any open-extend pair (*x, y*) from the original list with *a ≤ x ≤ c* and *b ≤ y ≤ d*).
2. We performed a search benchmark for the validation query set against all training and validation proteins for each combination of open and extend in the reduced list determined by the previous task. We computed the average sensitivity up to the first false positive at the family, superfamily, and fold levels. We then selected the gap open and extend parameters with the highest result at the superfamily level.

We followed a similar procedure for determining the optimal weighting when combining different alphabets. In the two alphabet case, we considered weights on the first alphabet of *{*0.3, 0.4, 0.5, 0.6, 0.7*}* and chose the second alphabet weight so the weights summed to one. In the three alphabet case we considered 15 weighting schemes in which each alphabet had weight at least 0.2 and all weights are of the form *x/*10 for an integer *x*. We performed the validation alignment task with all combinations of weights and gap open and extend parameters listed above. When the optimal parameters determined by mean lDDT and spearman correlation included the same weighting, we fixed this weighting and followed the procedure above to select the smaller list of gap open and extend parameters for the validation search benchmark. In cases where different weightings produced the best mean lDDT and spearman correlation, we considered all combinations of weightings and gap parameters that yielded mean lDDT or spearman correlations that were sufficiently close to the maximums.

Table A.2 gives the parameters used for each method.

### A.3 Other discretization approaches

#### A.3.1 VQVAE

Following the success of Foldseek in using a VQ-VAE as an effective clustering tool, we adapted the technique to our *n*-hot data as follows. The *encoder* maps each *n*-hot vector through three fully-connected layers to a continuous 2-dimensional latent representation. This latent representation mapped via the *vector quantizer* to the nearest (*l*_2_) centroid in a learned set of 20 centroids. The *decoder* maps the centroid associated with each input *x* to a length-1000 vector 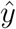 intended to predict *n*-hot vector of the amino acid aligned with *x*. We train the VQ-VAE on pairs of aligned positions using a Jaccard loss function (see Equation (1)).

We associate each of the 20 centroids with a letter in our alphabet. Given a *L ×* 1000 representation of a protein, we render a sequence of length *L* by running each length-1000 *n*-hot vector through the encoder and vector quantizer portions of the VQ-VAE. Each encoded, quantized input takes the form of a centroid, associated with some character in our 20-letter alphabet. We assign this character to the position. Note the decoder is used only in training.

Our VQ-VAE was implemented in PyTorch (version 2.4.1) over a custom Jaccard (IoU) loss function, with commitment cost = 0.25, the Adam optimizer, a batch size of 512, and a learning rate of 10^*−*3^ over six epochs. The encoder portion comprises layers of 1000, 1000, and 20 nodes; the decoder comprises layers of 20, 1000, and 1000 nodes.

Clusters found by VQ-VAE did not reach the quality of the clusters generated by our graph clustering method nor are they as interpretable. The sensitivities up to the first false positive at the family, superfamily, and fold levels were 0.756, 0.333, 0.053 respectively. This is similar to the performance of the dihedral alphabet at the family level and a slight improvement for the superfamily and fold levels.

#### A.3.2 Optimizing learned centers with mutual information

Our graph clustering approach learns 20 cluster centers and classifies each amino acid as the character corresponding to the closest cluster center with respect to the Jaccard metric. We attempted to further optimize the location of the cluster centers using gradient descent with an objective function that sought to jointly maximize the mutual information and entropy of a BLOSUM matrix computed for random subsets of the training data. After adjusting the cluster centers in this manner, the corresponding alphabet yielded sensitivities up to the first false positive at the family, superfamily, and fold levels of 0.804, 0.385, 0.077 respectively. This performance was marginally worse than our graph-cluster derived 3Dn alphabet.

### A.4 Details on scoring schemes

The following details the computation of the entries *M*_*ij*_ of similarity matrix between two proteins with the blurry neighborhood method. First we compute the Jaccard metric between the blurry neighborhoods of the *i*^*th*^ amino acid in the first protein and the *j*^*th*^ amino acid in the second protein. Then we apply a transformation that maps the value of Jaccard metric to a log-odds score. We compute this transformation as follows. We compute the Jaccard metric for pairs of aligned positions and non-aligned positions in the the training set. We bin the Jaccard values into 100 bins (e.g. 0-0.01; 0.01-0.02; … 0.99-1.0). For each bin, we take twice the *log* of the ratio of the number of aligned pairs with Jaccard in that bin to the total number of pairs with Jaccard in that bin. This value is the transformed Jaccard score for the bin. We manually adjust the values for high and low Jaccard bins with little data.

For our alphabets, the entry *M*_*ij*_ is given by a BLOSUM score for the *i*^*th*^ character of the first sequence and the *j*^*th*^ character of the second sequence. Our BLOSUM scores are computed as follows:

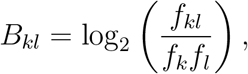

where *f*_*kl*_ is the the observed frequency of alignment of characters *k* and *l*, and *f*_*k*_ and *f*_*l*_ are the observed frequency of characters *k* and *l*, respectively.

### A.5 Dihedral BLOSUM

Here we describe the procedure to construct an alphabet using dihedral angles. We start by computing the backbone dihedral angles *ϕ* and *ψ* for proteins in our dataset. The distribution of these angles (points on the Ramachandran plot) is binned into 30 *×* 30 bins (bins of size 12^*°*^ *×* 12^*°*^) to identify high-frequency bins that each account for at least a threshold fraction (*>* 1*/*900) of total number of points. We use the same threshold separately for *PRO* and *GLY* due to their propensity to occupy non-standard positions in the Ramachandran plot. Points that are not in these high-frequency bins are discarded and we only focus on these high-frequency bins for downstream analyses. Thus, the procedure reduces the total number of bins from 900 to 251.

Next, we construct a 251 *×* 251 substitution probability matrix *P* that expresses how often two bins substitute for each other among structurally aligned protein pairs. An element in the matrix is given by 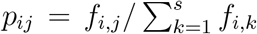, where *f*_*i,j*_ is the number of times bin *i* substitutes for bin *j* and *s* is the alphabet size. The denominator counts the total number of times bin *i* substitutes for any bin.

Then we reduce the number of bins by performing a clustering algorithm on the probability matrix that seeks to merge bins in a way that maximizes the mutual information of the distribution. The mutual information of a probability matrix *P* is *MI* =∑ _*i,j*_ *p*_*ij*_ log(*p*_*ij*_)*/*(*p*_*i*_*p*_*j*_). We merge a pair of bins results in the maximum *MI* over the new set of *n −* 1 bins. The idea is to have a small number of bins without significant decrease in *MI*. When attempting to merge a pair of bins, we only consider the bins that are adjacent with periodic boundary condition. We continue merging bins until no further bins can be merged, i.e. no remaining bin pairs are contiguous. Our goal is to construct a 20-character alphabet for a fair comparison with other methods. We select 18 as the appropriate number of hyperbins (clusters of original 251 bins, labeled 0 - 17). Two additional characters are included in the alphabet: one for endpoints of the sequences where the dihedral angles cannot be computed (18), and the other for angles that fall outside the high-frequency bins (19). The structure of a protein can thus be represented as a sequence of these dihedral cluster labels (similar to amino acid sequence), which we use to construct the dihedral BLOSUM matrix.

### A.6 Description of data

For the database search task, we utilized a dataset of 11211 proteins. Folds from SCOPe40 [3] were split 80%/20% into training and test sets, with 10% of the training set reserved for validation. The transition matrix and dihedral BLOSUM clusters were training utilizing a set of TM-alignments of protein pairs with TM-scores of at least 0.6. We used the validation set to train algorithm hyperparameters: the open and extend gap penalty for the Smith-Waterman algorithm and the relative weights of the alphabet combinations. The process detailing the hyperparameter selection process is outlined in Appendix A.2. When executing the search benchmarking task, we queried each test protein against the entire protein database.

## B Supplemental Figures

## Footnotes

1 Inferred beta carbon in the case of glycine.

